# A *Caenorhabditis elegans* model of adenylosuccinate lyase deficiency reveals neuromuscular and reproductive phenotypes of distinct etiology

**DOI:** 10.1101/181719

**Authors:** Adam R. Fenton, Haley N. Janowitz, Melanie R. McReynolds, Wenqing Wang, Wendy Hanna-Rose

## Abstract

Inborn errors of purine metabolism are rare syndromes with an array of complex phenotypes in humans. One such disorder, adenylosuccinate lyase deficiency (ASLD), is caused by a decrease in the activity of the bi-functional purine biosynthetic enzyme, adenylosuccinate lyase (ADSL). Mutations in human ADSL cause epilepsy, muscle ataxia, and autistic-like symptoms. Although the genetic basis of ASLD syndrome is known, the molecular mechanisms driving phenotypic outcome are not. Here, we characterize neuromuscular and reproductive phenotypes associated with a deficiency of *adsl-1* in *Caenorhabditis elegans.* Characterization of the neuromuscular phenotype reveals a disruption of cholinergic transmission affecting muscular contraction. Using genetics, pharmacological supplementation, and metabolite measurements, we correlate phenotypes with distinct metabolic perturbations. The neuromuscular defect is associated with a toxic accumulation of a purine biosynthetic intermediate whereas the reproductive defect can be ameliorated by purine supplementation, indicating differing molecular mechanisms behind the phenotypes of ASLD. Because purine metabolism is highly conserved in metazoans, we suggest that similar separable metabolic perturbations result in the varied symptoms in the human disorder and that a dual-approach therapeutic strategy may be beneficial.

## Author summary

Adenylosuccinate lyase deficiency is a rare metabolic disorder that is associated with epilepsy, muscle ataxia, and autistic-like symptoms in humans. This disorder arises from mutations in adenylosuccinate lyase, an enzyme involved in purine nucleotide biosynthesis. While we understand the genetic basis of this disorder, the mechanism of pathogenesis is unknown. Moreover, the linkage between phenotype and metabolic perturbation remains unclear. We report here on neuromuscular and reproductive phenotypes caused by a deficiency of *adsl-1* in *Caenorhabditis elegans.* For each defect, we identified a specific metabolic perturbation that causes the phenotype. The neuromuscular phenotype is associated with a toxic accumulation of a purine metabolic intermediate whereas the reproductive phenotype can be alleviated by purine supplementation. Our results point to separate molecular mechanisms as causative for the phenotypes, suggesting that there may be a similar relationship between phenotype and metabolic perturbation in humans. As such, our model suggests the use of a multi-pronged approach in humans to therapeutically target the metabolic perturbation contributing to each symptom.

## Introduction

Inborn errors of purine metabolism are understudied syndromes that arise from mutation of purine biosynthetic or catabolic enzymes. Although rare, these disorders are thought to be underdiagnosed because the varied clinical symptoms mimic other disorders (1). Purine disorders can have devastating health effects and often result in early death. Not only are there few therapeutic options available to patients, but the intriguing biological mechanisms linking defects in purine biosynthesis to phenotypic outcomes have also been difficult to decipher. Our aim is to use a fast, inexpensive and yet applicable model to explore the molecular links between perturbations in purine biosynthesis and organismal physiological and behavioral outcomes and to generate therapeutic strategies for these rare and understudied syndromes.

Purine nucleotides are monomers that polymerize with pyrimidine nucleotides to form nucleic acids. They also serve critical roles in cell signaling, energy storage and transfer, and metabolic regulation (2). Purines are synthesized via two biosynthetic pathways: *de novo* and salvage. *De novo* purine biosynthesis forms purine monomers from the components of intracellular amino acids and sugars. This pathway takes eleven steps to convert ribose-5-phosphate (R5P) to inosine monophosphate (IMP), the precursor for other purine monomers (Fig 1). The salvage biosynthetic pathway uses nucleic acid constituents from the diet or purine catabolism to create new purine products.

**Fig 1.**
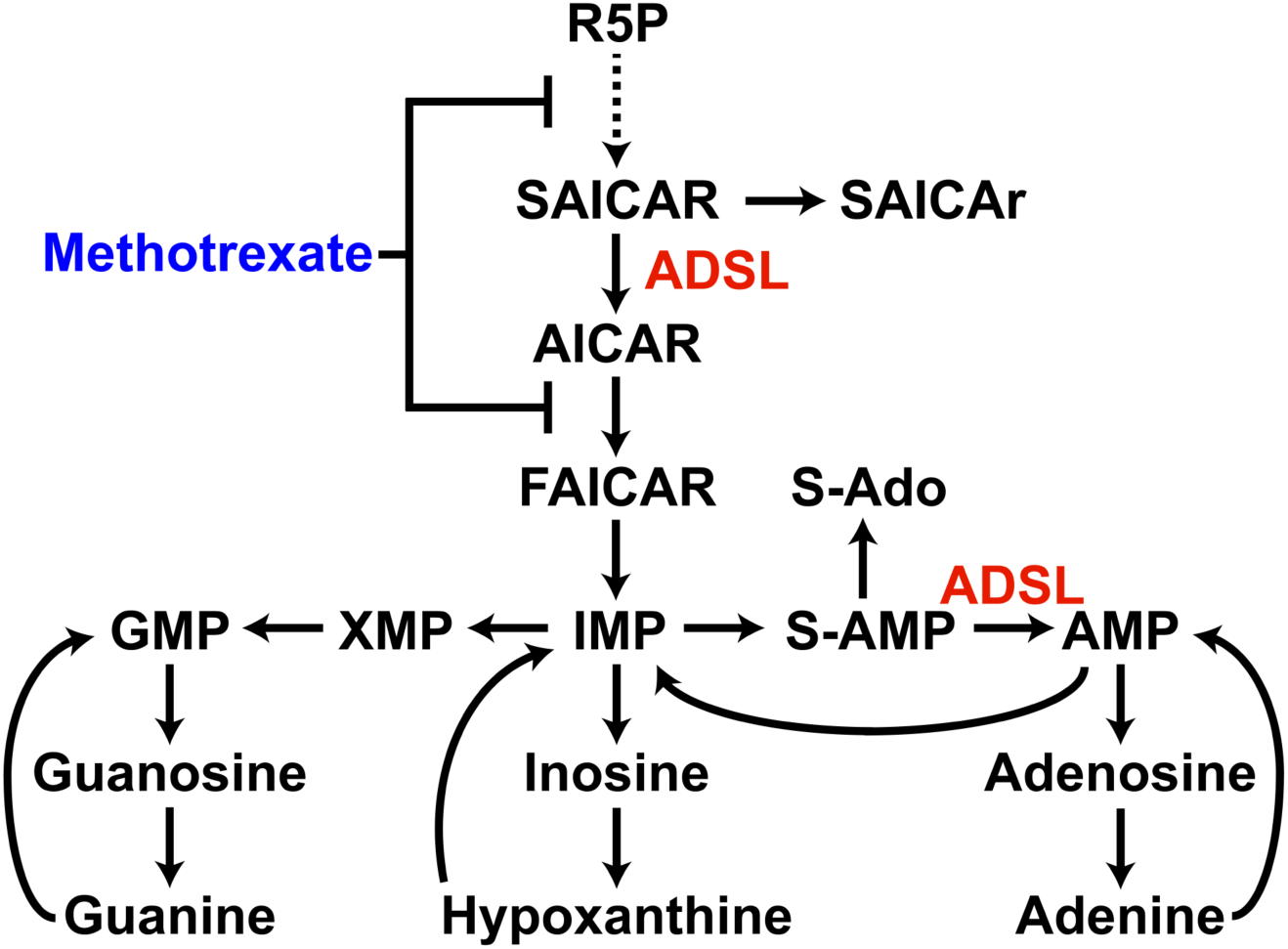
Purine biosynthesis pathways. ADSL functions twice in *de* novo purine biosynthesis. Methotrexate is an antimetabolite that indirectly inhibits *de novo* synthesis. Abbreviations: R5P, ribose-5-phospate; SAICAR, succinylaminoimidazole carboxamide ribotide; SAICAr, succinylaminoimidazole carboxamide riboside; ADSL, adenylosuccinate lyase; AICAR, aminoimidazole carboxamide ribotide; IMP, inosine monophosphate; S-AMP, adenylosuccinate; S-Ado, succinyladenosine AMP, adenosine monophosphate; XMP, xanthine monophosphate; GMP, guanosine monophosphate.

Adenylosuccinate lyase (ADSL) is an enzyme with dual functions in *de novo* purine biosynthesis. It catalyzes the cleavage of succinyl groups to yield fumarate twice in *de novo* synthesis; it converts succinylaminoimidazole carboxamide ribotide (SAICAR) to aminoimidazole carboxamide ribotide (AICAR) and succinyladenosine monophosphate (S-AMP) to adenosine monophosphate (AMP). Adenylosuccinate lyase deficiency (ASLD) is a human syndrome associated with a spectrum of symptoms including seizures, ataxia, cognitive impairment, and autistic-like behaviors (3–5). Symptoms range in severity from mild to severe and are negatively correlated with the degree of residual ADSL activity (6). In the most extreme cases, ASLD is neonatally fatal due to prenatal growth restriction, encephalopathy, and intractable seizures (6,7). This autosomal recessive neurometabolic disorder has been reported in over 50 patients since its original characterization in 1969; for these cases, over 40 separate mutations in adenylosuccinate lyase (ADSL) are associated with the disease state (8–10).

There are competing hypotheses about the etiology of ASLD symptoms. Severity of symptoms has been positively correlated with the level of accumulation of two succinylnucleosides, SAICAr and S-Ado, in the urine and cerebrospinal fluid (9,11). These nucleosides are the dephosphorylated forms of the ADSL substrates SAICAR and S-AMP, respectively, and their accumulation in body fluids is the only biochemical marker of the disorder. Previous findings associated a lower ratio of S-Ado/SAICAr with more severe symptoms, and it was hypothesized that S-Ado is protective while SAICAr is toxic (9,11). Recent findings indicate that this ratio is not predictive of phenotype severity, but correlates to the patient’s development and age during sample collection (6). Dephosphorylation of SAICAR to SAICAr has also been proposed to be a detoxification mechanism to reduce the toxic accumulation of SAICAR in affected cells (12). Thus, questions remain about the role of ADSL substrates in disease etiology.

It is also hypothesized the blockage of purine biosynthesis specifically contributes to ASLD symptoms. Deficiency of ADSL is expected to result in decreased concentrations of purine products, particularly adenine nucleotides, due to the dual function of this enzyme in the biosynthesis of AMP. However, no deficit in purines has been detected in patients; measurements of purine levels in kidney, liver, and muscle cells of ASLD patients are normal (13). Residual activity in patients likely contributes to the conservation of purine levels. Measurements of ADSL enzyme activity indicate that 3% residual activity is sufficient to convert S-AMP to AMP; although metabolic flux is greatly hindered (13). Moreover, a reduction in ADSL activity can be circumvented via supply of purines through the salvage pathway and dietary intake (14). In this case, affected cells and tissues would be dependent on high activities of the salvage enzymes to maintain purine levels. It remains possible that a deficit in the ability to synthesize purines *de novo* at a specific developmental stage contributes to phenotypic outcome, but evidence in support of these hypotheses to explain ADSL phenotypes is still lacking.

The pathological mechanisms causing the disorder also remain unknown (15–17). We are interested in probing the mechanism behind the disorder using *Caenorhabditis elegans*, an established organism for studying metabolism and associated metabolic disorders (18). The purine metabolic pathways are highly conserved across all eukaryotes, including *C. elegans* (19). This level of conservation indicates the functionality of *C. elegans* as a model for errors of purine metabolism. In addition to metabolic conservation, *C. elegans* provides a well-characterized nervous system that is essential for studying symptomatic aspects of ASLD. By using a model with a simple and fully identified neural network (20), the nervous system function can be studied under conditions of ADSL depletion. Thus, *C. elegans* has physiological benefits that other models, such as mammalian cell culture and yeast (21,22), do not provide.

We report here on the development of *C. elegans* as a model for ASLD using a mutant allele and RNAi knockdown of the *adsl-1* gene. Extensive analysis of locomotive and reproductive phenotypes gives insight into which biological processes are disrupted by a decrease in ADSL function. We find that altered cholinergic synaptic transmission impacts muscle function in mutants. We examine metabolite levels in control and *adsl-1* animals and use pharmacological and metabolite supplementation to associate substrate accumulation and purine production with the different phenotypes of ASLD in *C. elegans.* We propose a similar linkage between metabolic perturbation and phenotype in humans due to the high level of conservation of *de novo* purine biosynthesis.

## Results

We used the *adsl-1(tm3328)* mutant and RNAi of *adsl-1* in the RNAi hypersensitive strain *eri-1(mg366)* (23,24) to model adenolyosuccinate lyase deficiency. The *tm3328* allele is a 792 bp deletion that removes over half of the *adsl-1* coding sequence, including the N-terminus. RNAi of *adsl-1* results in efficient yet incomplete knockdown of message levels to approximately 20% of controls (Fig 2A). We observed reproductive, developmental and locomotion defects in both *adsl-1(tm3328)* and *adsl-1(RNAi)* animals.

**Fig 2.**
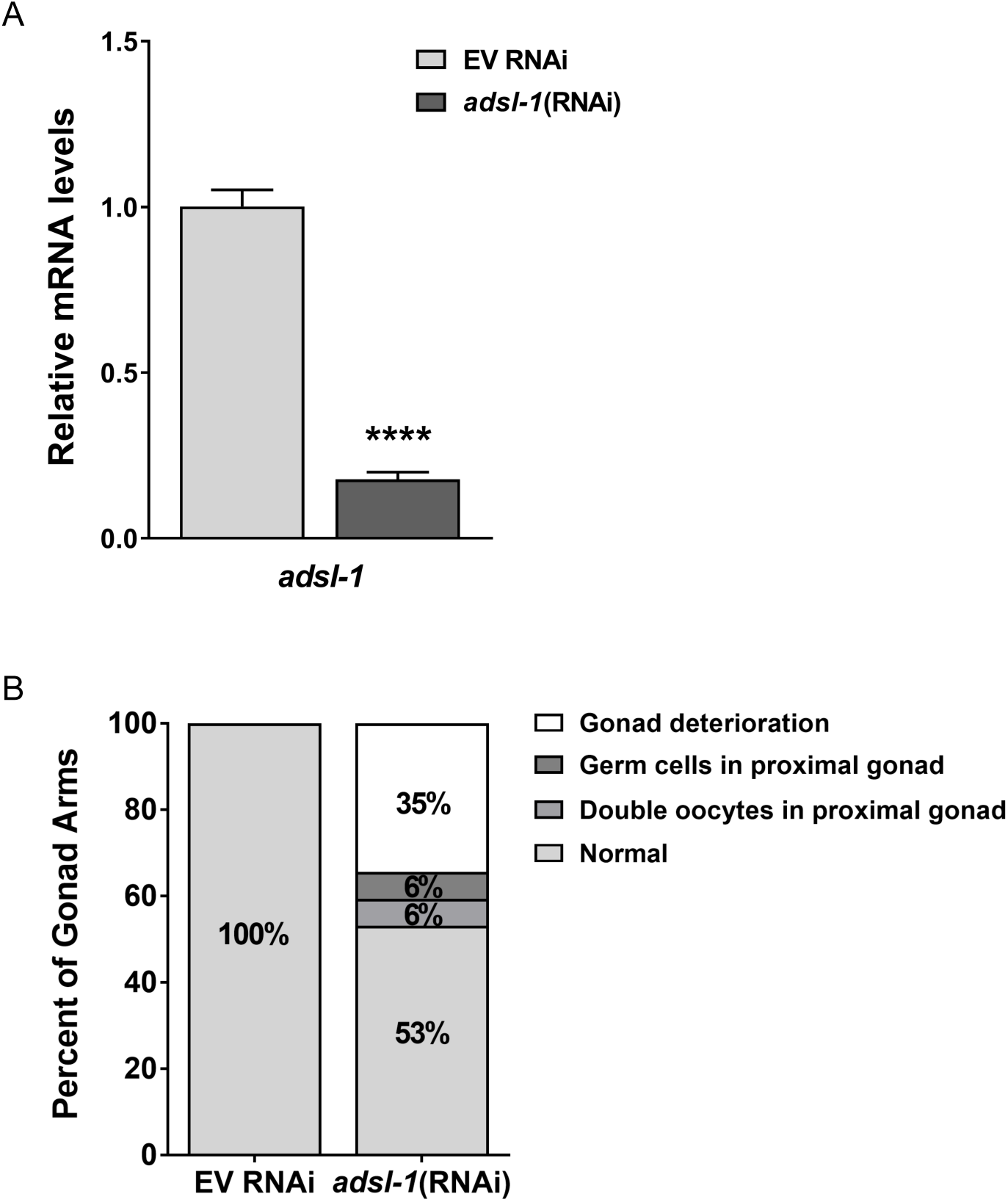
Disruption of *adsl-1* function causes reproductive defects. **(A)** *eri-1* animals were exposed to empty RNAi vector (EV) or *adsl-1*(RNAi) beginning at the mid-fourth larval stage and qRT-PCR was used to measure relative mRNA levels of *adsl-1* in subsequent generation. Values are averages from two biological replicates, performed in duplicate. Error bars represent S.D. **** represents p<0.0001 by student’s two-tailed t test. **(B)** Gonad and oocyte development are disrupted within 24 hours of exposure of fertile first-day adults to *adsl-1*(RNAi). n = 32 animals per condition. Numbers inside bars indicate actual percentages.

### Disruption of *adsl-1* function results in reproductive defects and embryonic lethality

Neither *adsl-1(tm3328)* mutants nor animals exposed to RNAi of *adsl-1* for their whole life cycle are capable of producing progeny (n>100). Compared to N2 strain control animals, the gonad arms of *adsl-1(tm3328)* adults appear deformed, severely shrunken, and lack any indication of mature germ cell production (S1 Fig), indicating a requirement for *adsl-1* in normal gonadogenesis. To reveal processes that may require *adsl-1* function acutely, we also exposed fertile egg-laying adult animals in their first day of egg-laying to RNAi. Within 24 hours, these animals display an array of phenotypes. We observed both germ cells in the proximal gonad and oocytes in double file as opposed to single file in the proximal gonad, indicating abnormal progression of oogenesis (Fig 2). Deterioration of gonad arms was also evident (Fig 2, S2 Fig). We conclude that *adsl-1* is required for normal development of the gonad and is required acutely for maintenance of normal oogenesis.

Animals exposed to *adsl-1(*RNAi) starting in the mid-fourth larval stage produce early offspring that can be phenotypically examined. We observed a high degree of embryonic lethality (18%) in these offspring (Fig 3A). Thus, not only is oogenesis hindered when *adsl-1* function is decreased, but embryonic development is disrupted as well. We also examined the *adsl-1(tm3328)* mutant strain for evidence of developmental lethality. The sterility of the *adsl-1(tm3328)* strain requires the strain to be maintained using a balancer chromosome. Because the hT2 balancer is homozygous lethal, a genotypic ratio of one *adsl-1(tm3228)* homozygote for every two balanced heterozygotes should segregate from the balanced heterozygote. However, only 16% of the progeny of balanced heterozygotes were homozygous *adsl-1(tm3328/tm3328)* mutants (Fig 3B). This altered genotypic ratio of one homozygote for every 5.3 heterozygotes indicates that 62% of the *adsl-1(tm3328/tm3328)* population is missing. We conclude that there is a developmental lethality for the homozygous mutants, similar to the embryonic lethality of *adsl-1*(RNAi).

**Fig 3.**
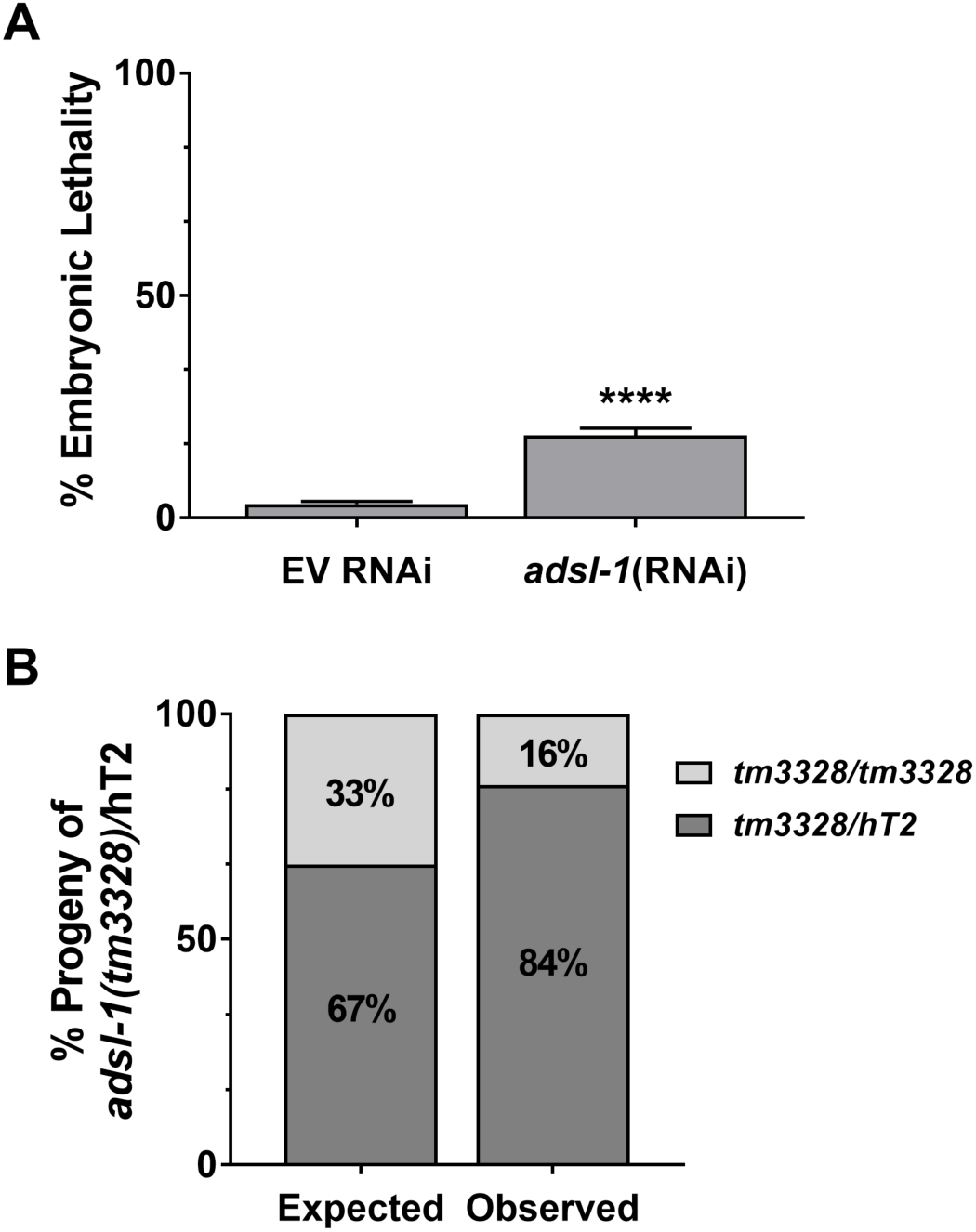
Disruption of *adsl-1* function causes developmental lethality. **(A)** Offspring of *eri-1* animals exposed to *adsl-1*(RNAi) beginning at the fourth larval stage display a high degree (18%) of embryonic lethality compared to the EV control (3%). n>450 eggs for each condition. Error bars are 95% confidence intervals. **** represents p<0.0001 using student’s two-tailed t test. (**B**) Homozygous *adsl-1(tm3328)* mutants are less prevalent than expected in the population of progeny that segregate from the balanced *adsl-1(tm3328)/*hT2 strain. n=695 animals for the observed genotypic ratio.

### Disruption of *adsl-1* function results in neuromuscular defects

*adsl-1(tm3328)* and *adsl-1*(RNAi) animals are noticeably sluggish compared to control animals. We quantified crawling speed of *adsl-1(tm3328)*, demonstrating that they have severely slowed locomotion (Fig 4A). Upon transfer to liquid, *C. elegans* will continually thrash for over 90 minutes before alternating to periods of inactivity (25). We also manually counted the thrashing rate during this active period as an indication of body wall muscle function. Thrashing rate is reduced for both *adsl-1(tm3328)* and *adsl-1*(RNAi) animals; mutants and RNAi animals exhibit a 77% and 22% reduction in thrashing rate, respectively (Fig 4B). The decreased phenotypic severity of *adsl-1*(RNAi) likely reflects the incomplete knockdown by RNAi (Fig 2A).

**Fig 4.**
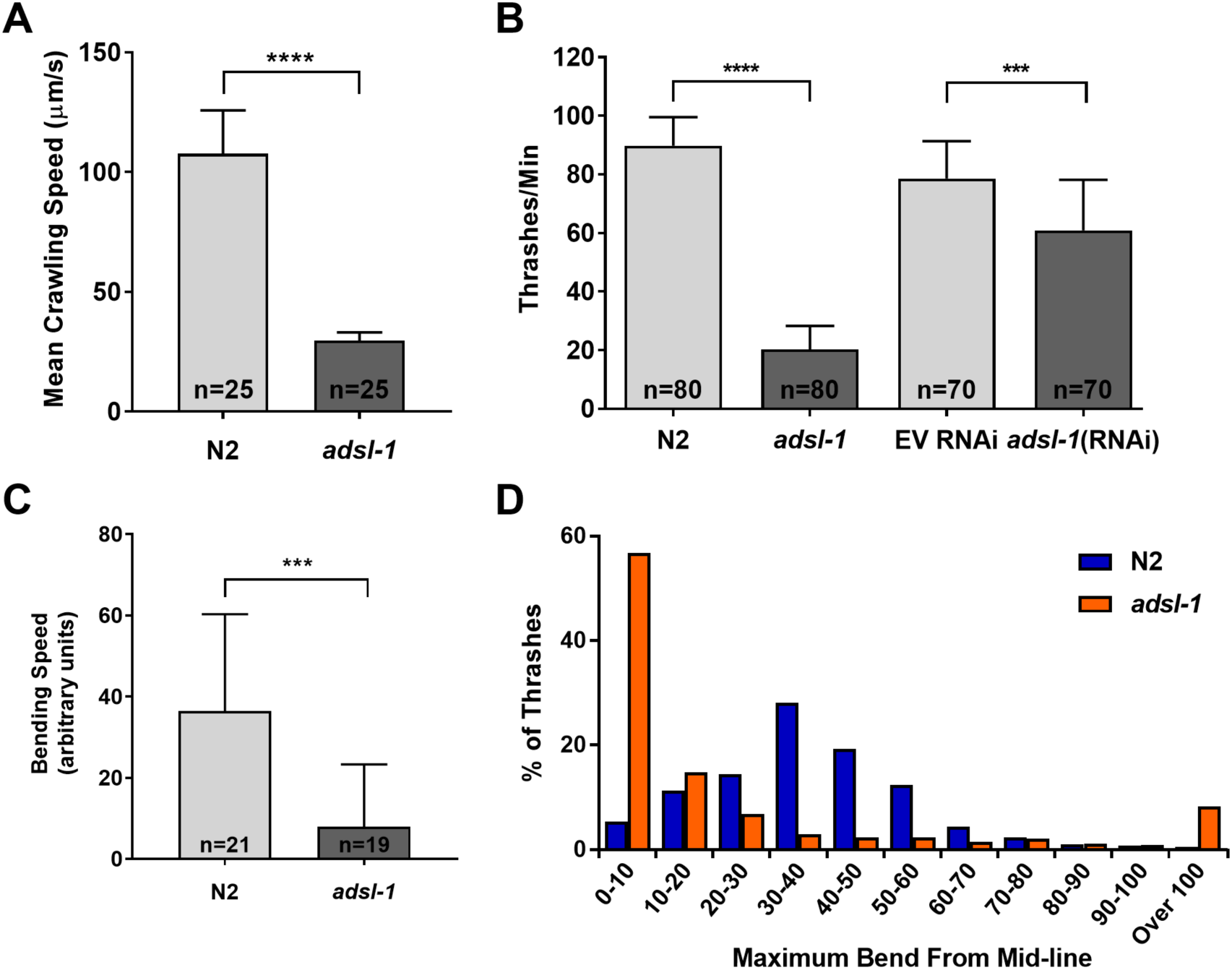
Disruption of ADSL function causes movement defects. (**A**) Comparison of average crawling speed for N2 control and *adsl-1(tm3328)* mutants. (**B**) *adsl-1(tm3328)* mutants and *adsl-*1(RNAi) have significantly decreased thrashing rate compared to N2 and empty vector (EV) control, respectively. (**C**) Average bending speed during thrashing reflects the decreased thrashing rate of *adsl-1(tm3328)* mutants. (**D**) N2 animals display a normal distribution for degree of bending while thrashing in liquid. *adsl-1(tm3328)* mutants primarily exhibit a smaller degree of bending during thrashing, but a proportion of bending angles are significantly more pronounced. Actual sample sizes indicated on each bar for A-C. In D, n = 20 animals per condition. Error bars indicate S.D. ***, and **** represent p<0.001, and p<0.0001, respectively, calculated using student’s two-tailed t test.

*adsl-1(tm3328)* animals appear uncoordinated in addition to their sluggish movement. Thus, we measured additional parameters of thrashing animals using ImageJ. The *adsl-1(tm3328)* animals have a 78% reduction in the average speed at which their body bends, consistent with manual counts of thrashing rates (Fig 4C). Control N2 animals exhibit an undulatory pattern of locomotion (26,27) with a normally distributed angle of bending intensity around an average of 37.8 while *adsl-1(tm3328)* animals display a clearly distinct distribution of bend intensities (Fig 4D). Mutant animals bend with less intensity relative to N2 controls during the majority of contractions. Despite the preference for these small bends, *adsl-1(tm3328)* are capable of contractions comparable to and beyond that of N2; a small proportion of mutant bends exceed the typical range of bending for N2 controls. Body bends of minimal or maximal intensity in *adsl-1(tm3328)* deviate greatly from the undulatory bending required for coordinated movement in *C. elegans* (28,29). We conclude that both pace and quality of muscle contractions is altered in *adsl-1* mutant animals.

### Disruption of *adsl-1* function affects cholinergic signaling

We next considered the question of how disruption of *adsl-1* function might result in the observed muscle contraction phenotypes. We investigated the hypotheses that hindered locomotion was caused by either disruption of the cholinergic synaptic transmission, which is required for potentiating action potential firing in body wall muscle, or the reduced response of the muscle cells to this signal (30). We assessed the functionality of pre-synaptic neurons and post-synaptic muscle tissue in the neuromuscular junction of cholinergic body wall muscles using levamisole and aldicarb. Levamisole is a cholinergic receptor agonist that stimulates body wall muscles to the point of paralysis (31–33). Because levamisole only affects postsynaptic function, resistance to levamisole is indicative of altered function in the muscle. *adsl-1(tm3328)* displayed mild resistance to levamisole over a five hour period of exposure. Following 24 hours of continual exposure, no difference was observed between *adsl-1(tm3328)* and N2 controls (Fig 5A). Resistance to aldicarb has been shown for mutants in both pre-synaptic and post-synaptic tissue (34). *adsl-1(tm3328)* displayed a strong resistance to aldicarb over a five hour period of exposure (Fig 5B). Following 24 hours of continual exposure, 25% of the *adsl-1(tm3328)* animals resisted paralysis (Fig 5B). We conclude that *adsl-1* mutants exhibit a stronger resistance to aldicarb than levamisole. The mild resistance to levamisole may indicate that muscle response is suboptimal. However, the resistance to aldicarb indicates that neural transmission is significantly affected by loss of *adsl-1* activity.

**Fig 5.**
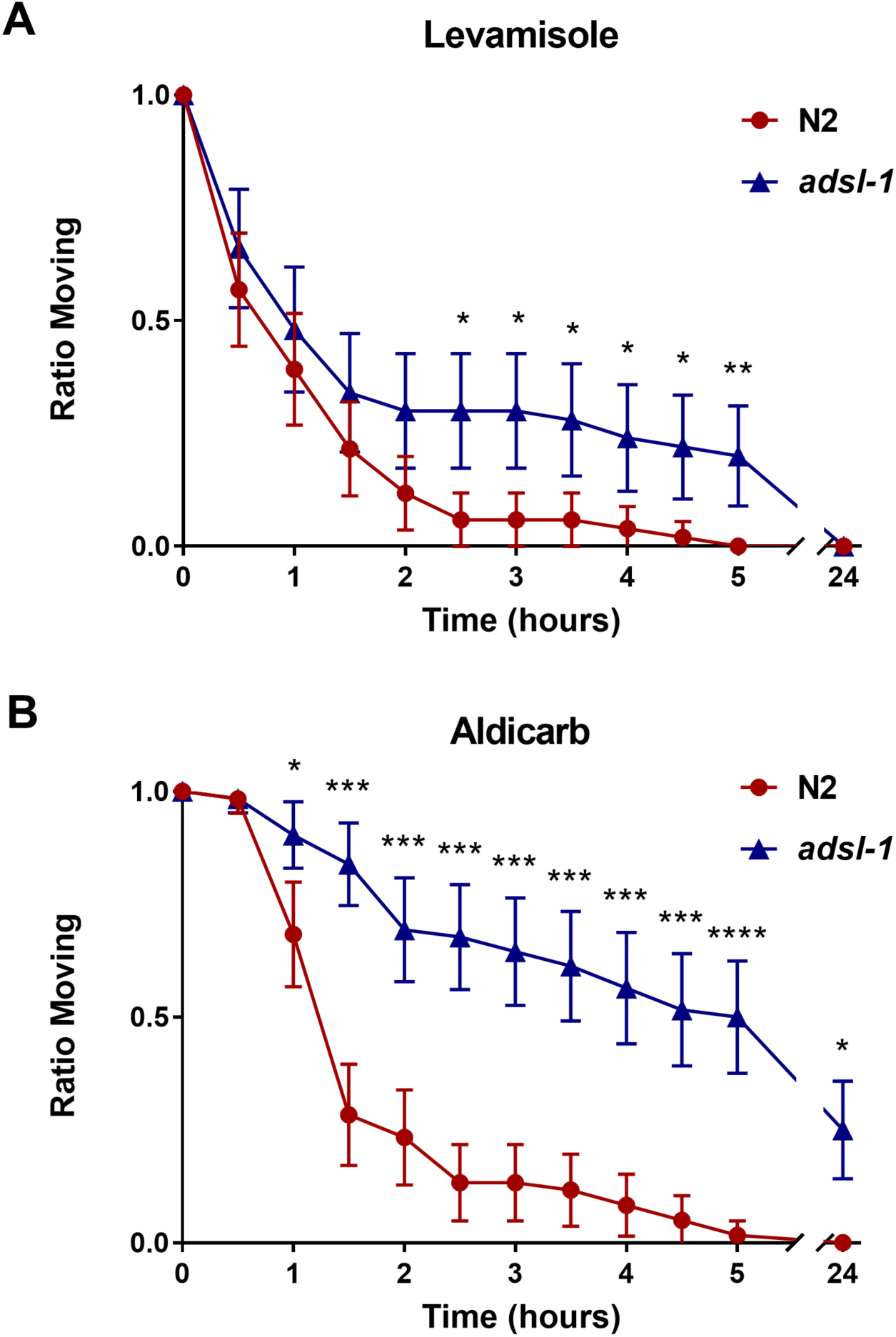
*adsl-1* mutants have altered cholinergic synaptic signaling. (**A**) *adsl-1* mutants are mildly resistant to the paralyzing effects of 1 mM levamisole. Resistance to levamisole is not sufficient to prevent paralysis after 24 hours of exposure. (**B**) *adsl-1* mutants are resistant to the paralyzing effects of 1 mM aldicarb. Complete resistance to aldicarb is displayed by a subpopulation of animals after 24 hours of exposure. Error bars are 95% confidence intervals. n = 50 – 62 animals per condition. *, 0.01 <p< 0.05; **, 0.001<p< 0.01; ***, p<0.001; ****, p<0.0001, calculated using two-tailed t test.

### Reduction of *adsl-1* function alters intermediate metabolite levels but has no effect on global purine levels

To investigate the hypotheses that changes in ADSL substrate or purine levels are causative of phenotypes, we quantified metabolite levels in *adsl-1*(RNAi) animals using LC-MS. We specifically measured the levels of both ADSL substrates, SAICAR and S-AMP, in six biological replicate samples of *adsl-*1(RNAi) and control *eri-1* animals. There were no detectable peaks for SAICAR in any of the control RNAi replicates (Fig 6A), indicating that the amount of SAICAR is typically below the threshold for metabolite detection via our methods. In all six replicates of *adsl-1*(RNAi), SAICAR was easily detected (Fig 6A), indicating that there is an increase in SAICAR levels when *adsl-1* is knocked down. Global levels of S-AMP are also increased in *adsl-1*(RNAi) compared to the control (Fig 6B). This data suggests that knockdown of *adsl-1* leads to the accumulation of ADSL substrates, similar to substrate accumulation shown in humans.

**Fig 6.**
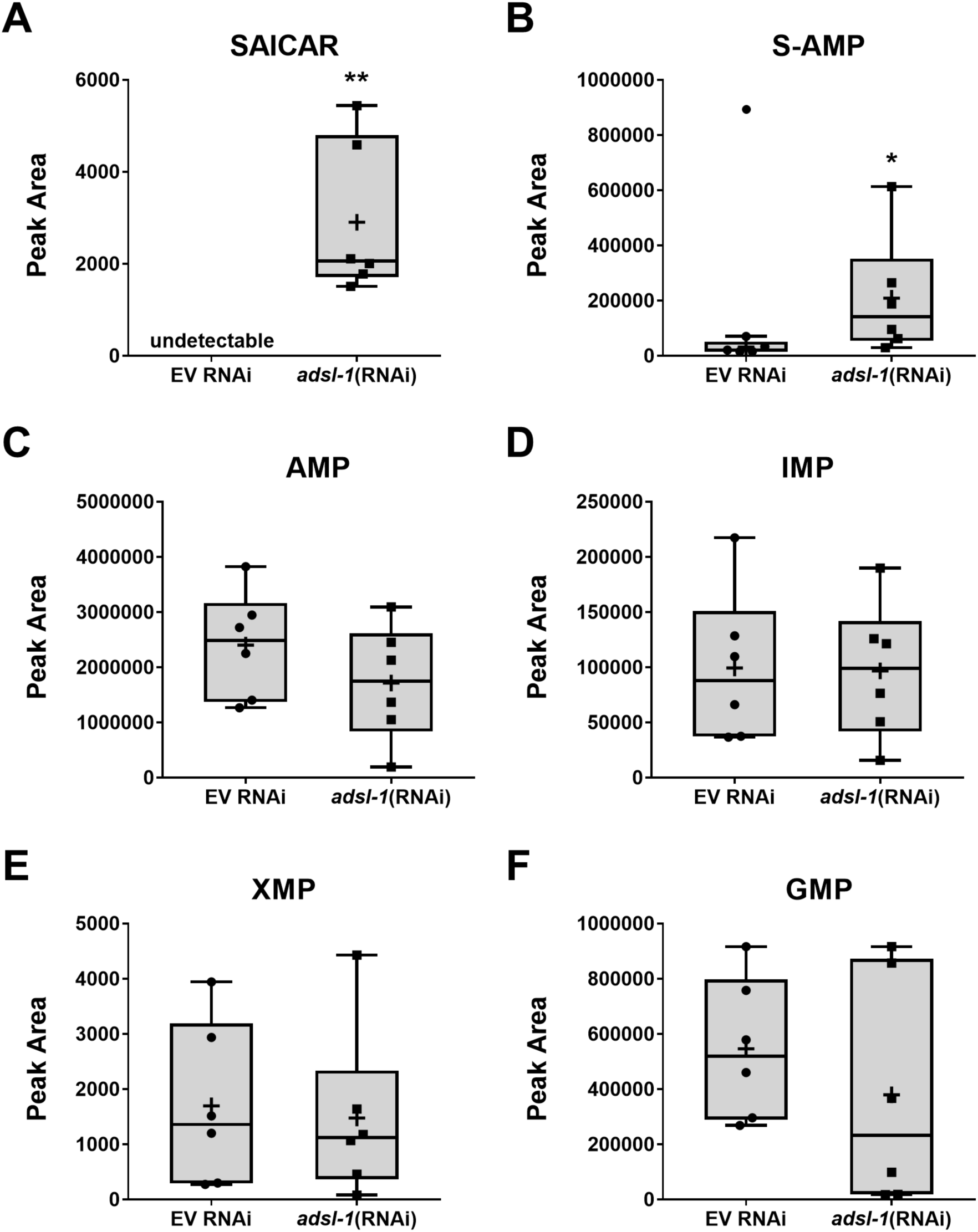
*adsl-1* knockdown causes substrate accumulation, but does not affect purine levels. (**A**) SAICAR is undetectable via LC-MS in the empty vector control and detectable for all biological replicates of *adsl-1*(RNAi). (**B**) S-AMP peak areas increase for *adsl-1*(RNAi) compared to the empty vector control. One of the data points for the empty vector control was a statistical outlier and not included in statistical analysis. (**C**) LC-MS peak areas for AMP trend downward for *adsl-1*(RNAi), but are not statistically significant from the empty vector control. (**D**) IMP peak areas are unchanged by RNAi knockdown of *adsl-1*. (**E**) LC-MS peak areas for XMP do not significantly differ for *adsl-1*(RNAi). (**F**) GMP peak areas are unchanged by RNAi knockdown of *adsl-1*. Each data point represents one biological replicate. Boxes show the upper and lower quartile values, + indicates the mean value, and lines indicate the median. Error bars indicate the maximum and minimum of the population distribution. *, 0.01<p<0.05; **, 0.001<p< 0.01, calculated using Welch’s t test.

We also measured global levels of purine monophosphate metabolites in *adsl-1(*RNAi) and control animals. Interestingly, none of these metabolites showed statistically significant changes upon *adsl-1* knockdown. AMP is the only metabolite that shows a downward trend (Fig 6C) with an average 37% decrease in the *adsl-1* samples, relative to controls. Levels of IMP do not exhibit any difference in *adsl-1*(RNAi) compared to the control (Fig 6D). Levels of XMP are more variable than that of AMP or IMP, but do not display any relative difference when comparing *adsl-*1(RNAi) to controls (Fig 6E). The relative levels of GMP have the largest variance of the examined metabolites for *adsl-1* RNAi, but did not exhibit a statistically significant difference from the control (Fig 6F). Overall, this data indicates that there is no significant decrease in purine metabolite levels caused by a knockdown of *adsl-1*.

### Reduced *de novo* synthesis contributes to the reproductive phenotype

To investigate the potential toxic effects of intermediate metabolite accumulation and the blockage of *de novo* purine production as causative of phenotypes, we examined the effect of both supplementation with purines and inhibition of substrate production on phenotypic outcome. Even though we detect no global deficit in purine levels, we investigated whether decreased purine production is functionally contributing to the reproductive phenotype by supplementing with purine products. This supplementation strategy would allow the purine salvage pathway to more efficiently compensate for the blockage of *de novo* biosynthesis. To block substrate accumulation, we used methotrexate, an antimetabolite that inhibits *de novo* purine biosynthesis upstream of ADSL (35,36).

Supplementation of cultures with purine products restored fertility in *adsl-1(*RNAi) animals. Fertility was restored to 90% of animals upon adenosine supplementation and 80% of animals upon guanosine supplementation (Fig 7A). Fecundity was also restored by supplementation with purines. Supplementation with adenosine restored fecundity to 65% of control levels and supplementation with guanosine restored fecundity to 62% of control levels (Fig 7B). Supplementation of cultures with methotrexate had no effect on the fecundity or fertility of *adsl-1* RNAi animals (Fig 7B); evidence for the uptake and inhibitory effect of methotrexate is shown below. Thus, the sterility phenotype is linked to a deficit in *de novo* purine synthesis, and we detected no role for substrate accumulation in the fertility phenotype.

**Fig 7.**
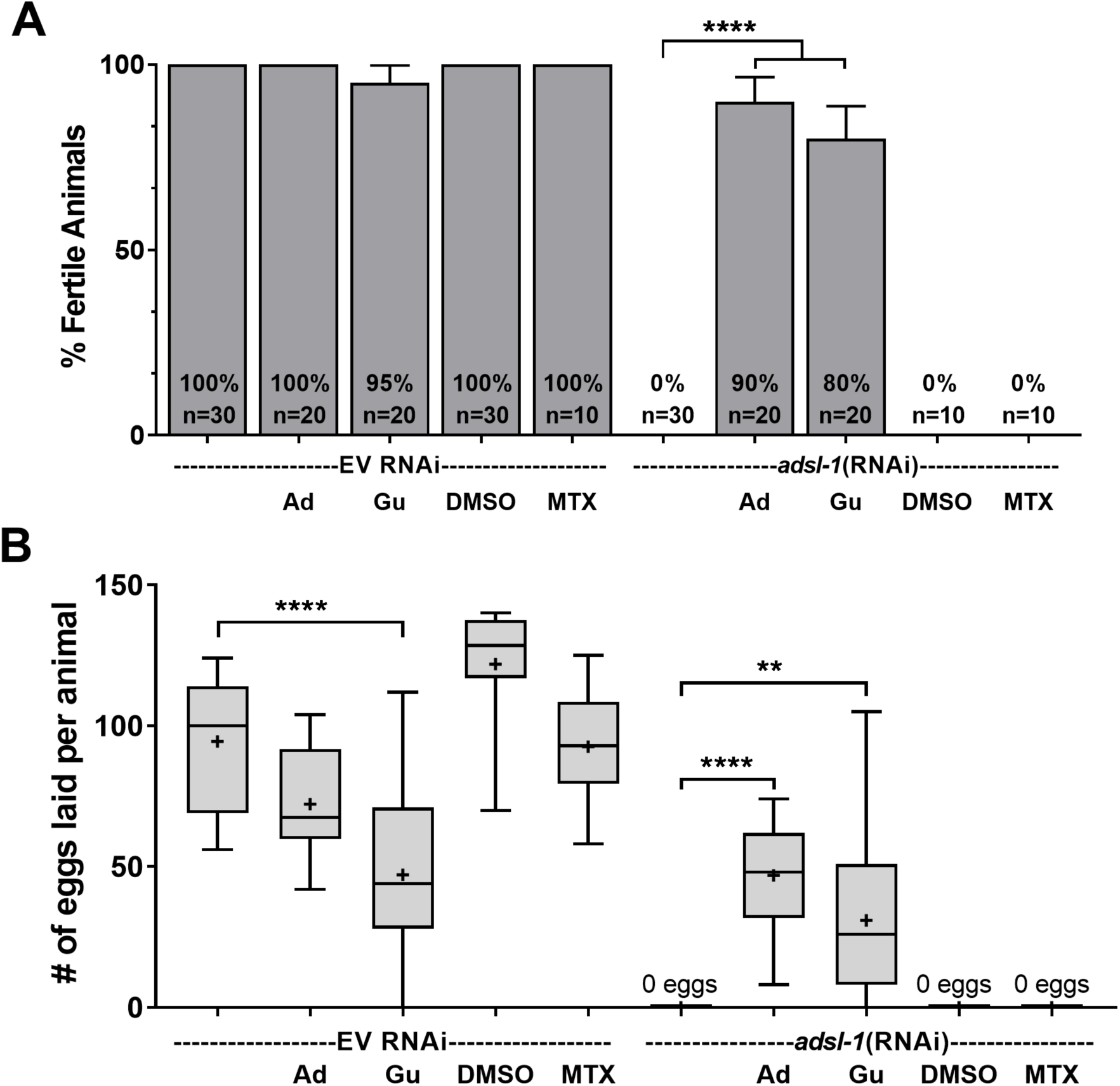
Purine supplementation restores fertility and fecundity to *adsl-1*. (**A**) Supplementation with 10 mM adenosine or guanosine restores fertility of *adsl-1*(RNAi) animals while supplementation with 22 μM methotrexate has no effect on fertility. (**B**) Supplementation with 10 mM adenosine or guanosine restores fecundity for *adsl-1*(RNAi). Supplementation with 10 μM methotrexate has no effect on fecundity in *adsl-1*(RNAi). In A, error bars are 95% confidence interval. Box plots are as described in Figure 6. n = 10-30 animals per condition. *, **, and **** represent p<0.05, p<0.01, and p<0.0001, respectively. Significance was calculated using ANOVA.

### Substrate buildup contributes to the neuromuscular phenotype

We also examined the effect of methotrexate and purine supplementation on the phenotypic outcome of *adsl-1(tm3328)* and *adsl-1*(RNAi) animals using thrashing assays. Both *adsl-1(tm3328)* and *adsl-1*(RNAi) displayed improved locomotion upon methotrexate supplementation. The supplemented mutants displayed a 212% increase in thrashing rate compared to the control mutants, but are only restored to ~45% of the N2 control (Fig 8A). The attenuation of the milder phenotype of *adsl-1*(RNAi) is more robust than that of the mutants; these animals thrash at a rate indistinguishable from the empty vector control (Fig 8B). We then used LC-MS to quantify the effects of methotrexate supplementation on *adsl-1*(RNAi) animals. As expected, methotrexate supplementation results in a decrease in SAICAR levels in *adsl-1*(RNAi) animals (Fig 8C). In contrast, the minor increase in S-AMP observed in *adsl-1*(RNAi) is not significantly affected by methotrexate supplementation (Fig 8D). Thus, methotrexate supplementation effectively decreases the accumulation of SAICAR, the first ADSL substrate in the de novo pathway. We also investigated whether a deficit in purine production is functionally contributing to the neuromuscular phenotype, similar to the reproductive phenotype. Supplementation with adenosine, sufficient to restore fertility and fecundity, had no effect on thrashing rate for *adsl-1(tm3328)* or the N2 control (Fig 8E). We conclude from these data that SAICAR accumulation likely affects neuromuscular function of *adsl-1*.

**Fig 8.**
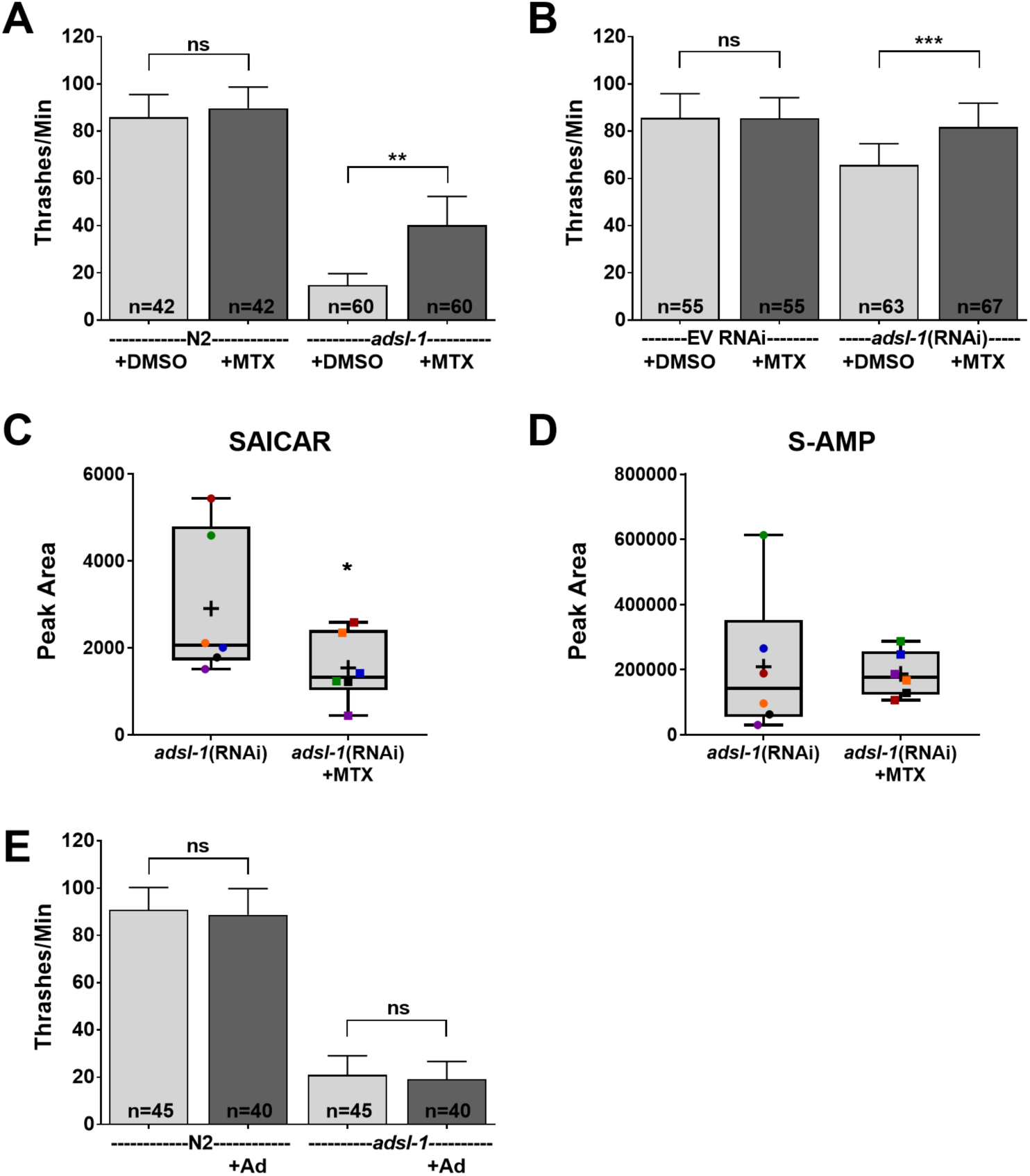
Upstream inhibition of de novo purine biosynthesis improves movement of *adsl-1*. (**A-B**) Supplementation with 22 μM methotrexate partially restores thrashing rate in *adsl-1* mutants (A) and fully restores thrashing rate in *adsl-1*(RNAi) (B). (**C**) LC-MS peak areas for SAICAR decrease in *adsl-1*(RNAi) upon supplementation with 22 μM methotrexate. (**D**) LC-MS peak areas for S-AMP do not change upon supplementation of *adsl-1*(RNAi) with methotrexate. (**E**) Supplementation with 10 mM adenosine has no effect *adsl-1* mutant thrashing rate. Colored data points indicate separate biological replicates of *adsl-1*(RNAi). Box plots are as described in Figure 5. Actual sample sizes indicated on each bar. Error bars indicate S.D. *, **, and *** represent p<0.05, p<0.01, and p<0.001, respectively. ns is not significant.

## Discussion

We have established *C. elegans* as an effective model for studying adenylosuccinate lyase deficiency (ASLD). *C. elegans* with reduced or eliminated function of ADSL have phenotypic and biochemical similarity to the human disorder. In both humans and *C. elegans*, individuals heterozygous for a mutation in ADSL are phenotypically normal, but homozygous individuals exhibit severe motor and developmental phenotypes (4,9,15). The locomotive defect in *C. elegans* mimics the muscle ataxia in human patients (15,16). Furthermore, metabolic analysis also shows similar substrate accumulation in whole animal lysates as in human patients (37). Phenotypic similarities were shown to be present in both *adsl-1* RNAi and *adsl-1(tm3328)* homozygotes, creating different genetic techniques to model this disorder.

Our observations regarding the sterility phenotype of *adsl-1* revealed disruption of both gonadogenesis and oogenesis. We found that the development of the gonad is severely hindered for both *adsl-1(tm3328)* and exposure to *adsl-1*(RNAi) during development. Additionally, normal oogenesis acutely requires the function of *adsl-1*. A decrease of *adsl-1* function following the L4 larval stage has minimal effect on gonadogenesis but disrupts the progression of maturing germ cells. Interestingly, the reproductive phenotypes are of similar severity when comparing the mutant to *adsl-1*(RNAi) animals. Given that the RNAi knockdown is incomplete, we conclude that the reproductive system is quite sensitive to the level of *adsl-1* activity for proper development.

Embryonic development is also sensitive to levels of *adsl-1* activity. We can avoid the typical sterility of *adsl-1*(RNAi) by administering the RNAi following larval development. Under these conditions, there is significant embryonic lethality associated with *adsl-1*(RNAi). We also demonstrated a developmental defect in the *adsl-1(tm3228)* mutants as evidenced by the deficit in homozygous mutants from the progeny of the balanced heterozygotes. However, this defect cannot be specifically linked to embryonic development. The fluorescent marker associated with the balancer is not visible in eggs, preventing selection of homozygous mutant eggs from the progeny pool. Embryonic lethality is likely to contribute to lethality of *adsl-1(tm3328)* as it does for *adsl-1*(RNAi), but the possibility remains that post-embryonic lethality or failure of mutant oocytes to mature contributes to the deficit of *tm3328* homozygotes in the balanced strain.

*adsl-1* animals are slow and display an irregular pattern of movement, mimicking the phenotypic outcome for ASLD in humans. Our data suggest a flaw in the muscle activation strategy behind the sinusoidal motion of *C. elegans* (38). While locomotion was slowed for both *adsl-1(tm3328)* and *adsl-1*(RNAi), the phenotype was more severe in *adsl-1(tm3328)* mutants. The milder locomotory phenotype of *adsl-1*(RNAi) likely reflects more residual enzyme activity and a less stringent requirement for high ADSL activity in locomotion compared to gonadogenesis and fertility. Nevertheless, *adsl-1*(RNAi) is an excellent model of the human syndrome, paralleling both a level of residual gene activity and a neuromuscular phenotype.

Given the evidence for a disruption in the patterning of muscle activation during locomotion, we investigated cholinergic signaling as a possible cause for this locomotive phenotype. A moderate resistance to levamisole in the *adsl-1* mutants indicates a variation in post-synaptic body wall function. Because levamisole can only stimulate the muscle tissue, we suggest that the resistance must arise from a defect in cholinergic receptors or the initiation of the contraction within the muscle itself. This assay reveals a tissue target that is involved in the locomotive phenotype. However, the more prominent resistance to aldicarb provides additional insight. Resistance to aldicarb can arise from defects in pre-synaptic acetylcholine release or from the post-synaptic cholinergic response. Because our levamisole assay exposed an issue with post-synaptic tissue, we examined the aldicarb resistance with this in mind. *adsl-1* mutants paralyze much slower on aldicarb and are capable of resisting paralysis past 24 hours of exposure. Because the aldicarb paralysis curve does not resemble that of levamisole, we conclude that there are additional factors contributing to the aldicarb resistance in pre-synaptic cholinergic neurons. This finding suggests that altered neuromuscular transmission may contribute to the muscular ataxia observed in ADSL patients.

By measuring metabolite levels for *adsl-1*(RNAi) animals, we have established that knockdown of *adsl-1* in *C. elegans* also results in metabolic similarity to the human syndrome. We did not measure metabolite levels in mutant strain because the sterility and developmental lethality associated with *adsl-1(tm3328)* prevents us from obtaining the large population of homozygous mutants required for LC-MS analysis. In addition to the limitations of the mutant strain, *adsl-1*(RNAi) was chosen for metabolomics analysis because this treatment is predicted to best model the human syndrome. The accumulation of the biosynthetic intermediates SAICAR and S-Ado during knockdown of *adsl-1* closely resembles the SAICAr and S-Ado accumulation observed across numerous human patients and tissue samples (3,6,11). As such, *adsl-1* knockdown in *C. elegans* mimics the primary diagnostic biochemical markers of ASLD in humans.

Interestingly, knockdown of *adsl-1* did not significantly affect any of the purine monophosphate products of *de novo* synthesis. The salvage biosynthesis pathways likely contribute to homeostatic mechanisms that maintain global purine levels in the absence of efficient *de novo* synthesis. It is likely that these animals are recycling enough purines from their diets to accommodate for the blockage of *de novo* biosynthesis but this model remains to be tested. Our metabolite measurements are derived from mixed-stage, whole-animal lysates. Thus, it remains possible that certain cells, tissues or developmental stages do not successfully maintain purine levels. Increased demand for purines or low activity of the salvage enzymes could alter purine levels for specific cells or developmental stages; more affected cell types could be masked by the whole-animal scale of metabolite measurements. Even with this possibility, the global maintenance of purine monophosphate levels still suggests compensation for the blockage of *de novo* synthesis in *adsl-1*(RNAi) animals. This maintenance of global purine levels is consistent with previous findings for adenine and guanine concentrations in patients and disease models with decreased ADSL function (13,21). Once again, metabolic profiling of *adsl-1*(RNAi) in *C. elegans* mimics the findings for human patients with decreased ADSL function, indicating the effectiveness of this model for studying ASLD.

Supplementation with individual purine products results in restoration of fertility. Each supplement can be converted to IMP, the central metabolite of purine synthesis, through the salvage pathways. In this way, these supplementations can overcome the blockage of IMP biosynthesis that results from the first function of ADSL, conversion of SAICAR to AICAR. However, adenosine is the only supplement that overcomes the second blockage of ADSL function, conversion of S-AMP to AMP. For this reason, it is interesting that both of the tested purine supplements are able to independently reverse the sterility of *adsl-1*(RNAi). We observed that adenosine supplementation is more robust than guanosine, but guanosine is still capable of restoring fertility to a significant extent. This result suggests that compensation for the second enzymatic function of ADSL is not as crucial for restoring fertility to *adsl-1*(RNAi) or that residual levels of ADSL-1 more easily suffice for this biochemical step.

Fertility restoration upon purine product supplementation indicates that a decrease in *de novo* purine production contributes to this phenotype. Furthermore, the correlation of sterility to the blockage of *de novo* synthesis is also predicted to be related to a potential increased demand for purines during the rapid division in gonad development when germ cells are dividing. A high demand for purines during reproductive development may cause a gonad-specific deficit of purines that is not reflected through metabolomics analysis for whole animal lysates of mixed age. Due to the severity of ASLD in humans, reproduction is not an option, so any direct correlation with the reproductive phenotype is unknown. Despite this, the linkage of a phenotype to a blockage of *de novo* purine formation in *C. elegans* indicates some of the human symptoms may have the same linkage. For this reason, one possible therapeutic approach to ASLD would be to supplement additional purines to the diets of affected individuals in combination with a block to purine biosynthesis.

Our finding that methotrexate supplementation alleviates the locomotive defect for both *adsl-1(tm3328)* and *adsl-1*(RNAi) suggests that substrate accumulation is causative of this phenotype. Metabolomics analysis of *adsl-1*(RNAi) specifically suggests that SAICAR accumulation is causative of this neuromuscular phenotype. Although methotrexate is capable of improving the locomotion of *adsl-1*(RNAi) to that of the empty vector control, methotrexate does not fully ameliorate the locomotive phenotype of *adsl-1(tm3328)*. It is possible that SAICAR accumulates to a greater extent in *adsl-1(tm3328)* than *adsl-1*(RNAi). In this case, methotrexate supplementation may not reduce SAICAR levels enough to fully attenuate the severe locomotive phenotype of *adsl-1(tm3328)*. Metabolomics analysis of *adsl-1(tm3328)* could reveal if this is the case, but is not technically feasible at this time.

The correlation between SAICAR accumulation and the locomotive defect is particularly interesting due to phenotypic similarity to muscular ataxia in humans. Because of this correlation, our data suggests that the motor control of humans may be improved by blocking SAICAR accumulation in a similar manner. The high conservation of purine biosynthesis indicates that a therapeutic approach using de novo synthesis inhibition could alleviate symptoms in humans. While these results expand on the relevance of SAICAR accumulation to phenotype, this study provides a crucial role in understanding the linkage between metabolic disturbance and disorder phenotype.

## Materials and methods

### *C. elegans* culture and strains

Strains were maintained on OP50 *Escherichia coli* as food under standard conditions at 20° C (39). We used the following strains; N2, *eri-1(mg366)*, and *adsl-1(tm3328).* The N2 and GR1373 *eri-1(mg366)* strains were obtained from the Caenorhabditis Genetics Center (CGC). *adsl-1(tm3328)* was obtained from the National BioResource Project in Tokyo, Japan and outcrossed three times against N2 The outcrossed, balanced strain was named HV854. This allele is homozygous sterile and was balanced with hT2, a balancer for the first and third chromosomes of *C. elegans*; this balancer causes pharyngeal expression of GFP (40). Non-GFP homozygous *adsl-1(tm3328)* animals were used in phenotypic analysis.

### RNAi

The *adsl-1* RNAi clone was from the *C. elegans* RNAi Library (Source BioScience, Nottingham, UK). RNAi feeding assays were carried out as described (23). Unless otherwise noted, we transferred mid-L4 *eri-1* animals to RNAi plates and examined their progeny in assays. *E. coli* strain HT115 carrying the empty RNAi feeding vector (EV) L4440 was used as a control.

### Metabolite Supplementation

We prepared filter-sterilized stock solutions of 22 mM methotrexate (Sigma) in DMSO and stock solutions of 117 mM adenosine (Sigma) in H_2_O with 10% 1 M NaOH and 150 mM guanosine (Sigma) in H_2_O with 25% 1 M NaOH. We added these solutions to OP50 seeded NGM plates to a final concentration of 22 μM methotrexate and 10 mM adenosine, guanosine. Following supplementation, we incubated the plates at room temperature for 1–2 days before use.

### Metabolomics

LC-MS metabolomics analysis was done with the Metabolomics Core Facility at Penn State. ~50 μL of animals were collected in ddH2O, flash frozen in liquid nitrogen and stored at 80°C. 15 μL samples were extracted in 1 mL of 3:3:2 acetonitrile:isopropanol:H2O with 1 μM chlorpropamide as internal standard. Samples were homogenized using a Precellys^(TM)^ 24 homogenizer. Extracts from samples were dried under vacuum, resuspended in HPLC Optima Water (Thermo Scientific) and divided into two fractions, one for LC-MS and one for BCA protein analysis. Samples were analyzed by LC-MS using a modified version of an ion pairing reversed phase negative ion electrospray ionization method (41).Samples were separated on a Supelco (Bellefonte, PA) Titan C18 column (100 x 2.1 mm 1.9 μm particle size) using a water-methanol gradient with tributylamine added to the aqueous mobile phase. The LC-MS platform consisted of Dionex Ultimate 3000 quaternary HPLC pump, 3000 column compartment, 3000 autosampler, and an Exactive plus orbitrap mass spectrometer controlled by Xcalibur 2.2 software (all from ThermoFisher Scientific, San Jose, CA). The HPLC column was maintained at 30°C and a flow rate of 200 uL/min. Solvent A was 3% aqueous methanol with 10 mM tributylamine and 15 mM acetic acid; solvent B was methanol. The gradient was 0 min., 0% B; 5 min., 20% B; 7.5 min., 20% B; 13 min., 55% B; 15.5 min., 95% B, 18.5 min., 95% B; 19 min., 0% B; 25 min 0% B. The orbitrap was operated in negative ion mode at maximum resolution (140,000) and scanned from m/z 85 to m/z 1000. Metabolite levels were corrected to protein concentrations determined by BCA assay (Thermo Fisher).

### Quantitative RT-PCR

Mid-L4 *eri-1* animals were placed on RNAi plates and RNA was isolated from mixed stage worms in the next generation using TRIZOL reagent (Invitrogen). 1 μg of RNA was converted to cDNA using the qScript cDNA Synthesis Kit (Quanta Biosciences). cDNA was diluted 1:10 and used for quantitative PCR using SYBR Green and Applied Biosciences RT-PCR machine. Three primer sets, *cdc-42*, *tba-1*, and *pmp-3*, were used as expression controls.

*cdc-42* F: ctgctggacaggaagattacg; R: ctgggacattctcgaatgaag

*tba-1* F: gtacactccactgatctctgctgaca; R: ctctgtacaagaggcaaacagccatg

*pmp-3* F: gttcccgtgttcatcactcat; R: acaccgtcgagaagctgtaga

*adsl-1* F: acagacaatggccgatcc; R: tgttggtttcaattccttggc

Results represent the average of two biological replicates each assayed in duplicate technical replicates.

### Phenotypic Analysis

#### Linear Crawling Velocity

Mid-L4 hermaphrodites were aged for 1 day at 20° C prior to the assay. Individual animals were tracked as they crawled on OP50 seeded NGM plates. 30 second videos were collected on a Nikon SMZ 1500 Stereoscope using NIS-Elements software from Nikon and analyzed using ImageJ. The mean linear crawling velocity was calculated for each animal by tracing the displacement of the animal’s midpoint. The displacement of the midpoint was tracked as a vector as the animal moved in a singular direction. Once the animal changed direction, a new vector was made to track movement in that direction; this process was repeated for the length of each video. Crawling velocity was determined by dividing each vector length by the corresponding time. The velocity values from all vectors in a video were averaged and adjusted for the time-fraction of each vector within the video.

#### Thrashing Assay

Mid-L4 hermaphrodites of each genotype were aged for 1 day at 20° C prior to the assay. Individual animals were placed in a drop of M9 solution on the surface of an unspotted NGM plate. After 1 minute of acclimation at room temperature, thrashes, the number of body bends, were counted for 1 minute using a Nikon SMZ645 Stereoscope.

#### Bending Quantification

Individual animals were aged and placed in M9 solution following the same procedure as the thrashing assay. 30 second videos were collected on a Nikon SMZ1500 Stereoscope using NIS-Elements software from Nikon. Videos were analyzed using ImageJ to create ideal conditions for computer-based quantification of *C. elegans* locomotion. First, the video background was subtracted using the rolling ball method. The background of each video is unchanging, allowing the starting frame to be subtracted from all frames. The videos were then converted to binary by setting a threshold with the “Otsu” thresholding algorithm. Binary videos of animals were processed through the wrMTrck plugin for ImageJ. Raw data of bending angle was obtained in the BendCalc format with bendDetect set to angle; this data provides bending angle for an animal at each frame of a video.

Raw data for the rate of change in bending angle were provided on a frame by frame basis for each video. The magnitude of these values were averaged to determine the bending speed for each animal; absolute magnitude was used to combine abduction and adduction into one dataset for all types of bending. Data points for the maximum bending extent were manually selected from raw data sets for each animal. The extent of each bend was determined by recording the value at each local maximum and minimum. These turning points represent the most extreme point in a bend before movement back to the mid-line of the animal. The absolute value of each bend was used for calculation of bending extent for each animal.

#### Paralysis

Two days prior to the assay, we added aldicarb (Sigma) or levamisole (Sigma) to unspotted NGM plates to a final concentration of 1 mM. We allowed the plates to dry at room temperature overnight then moved them to 4° C until the time of the assay. Approximately 20 mid-L4 hermaphrodites of each genotype were aged for 1 day at 20° C prior to the assay. We placed a 10 μl spot of OP50 *E. coli* solution on each plate and allowed it to dry for 30 minutes, concentrating the animals in a small area. Animals of a single genotype were placed on to an aldicarb plate and scored for paralysis every 30 minutes as described (34).

#### Egg-laying

For the egg-laying assay, ten *eri-*1 animals were placed on a plate containing the desired RNAi and supplementation. For each condition, ten second generation mid-L4 animals were placed onto individual plates. The number of eggs laid by each animal was counted over a five day period.

### Statistical Analysis

Two-tailed student t tests were used to determine p values when comparisons were limited between two conditions. One-way or two-way ANOVA was carried out with appropriate post-tests to determine p values between three or more experimental conditions. In LC-MS analysis, we used Welch’s two sample t test to calculate p values. We substituted all undetectable measurements with zeros to statistically compare conditions for LC-MS. In all figures: ns, not significant; *, 0.01 <p< 0.05; **, 0.001<p< 0.01; ***, p<0.001; ****, p<0.0001.

## Acknowledgements

We thank A. Patterson and P. Smith in the Penn State Metabolomics Core Facility for technical assistance and advice. *tm* alleles were provided by the Mitani laboratory through the National Bio-Resource Project of the Ministry of Education, Culture, Sports, Science, and Technology of Japan, Japan. Other strains were provided by the Caenorhabditis Genetics Center, which is funded by National Institutes of Health Office of Research Infrastructure Programs (Grant P40 OD010440).

## Supporting information

**S1 Fig.**
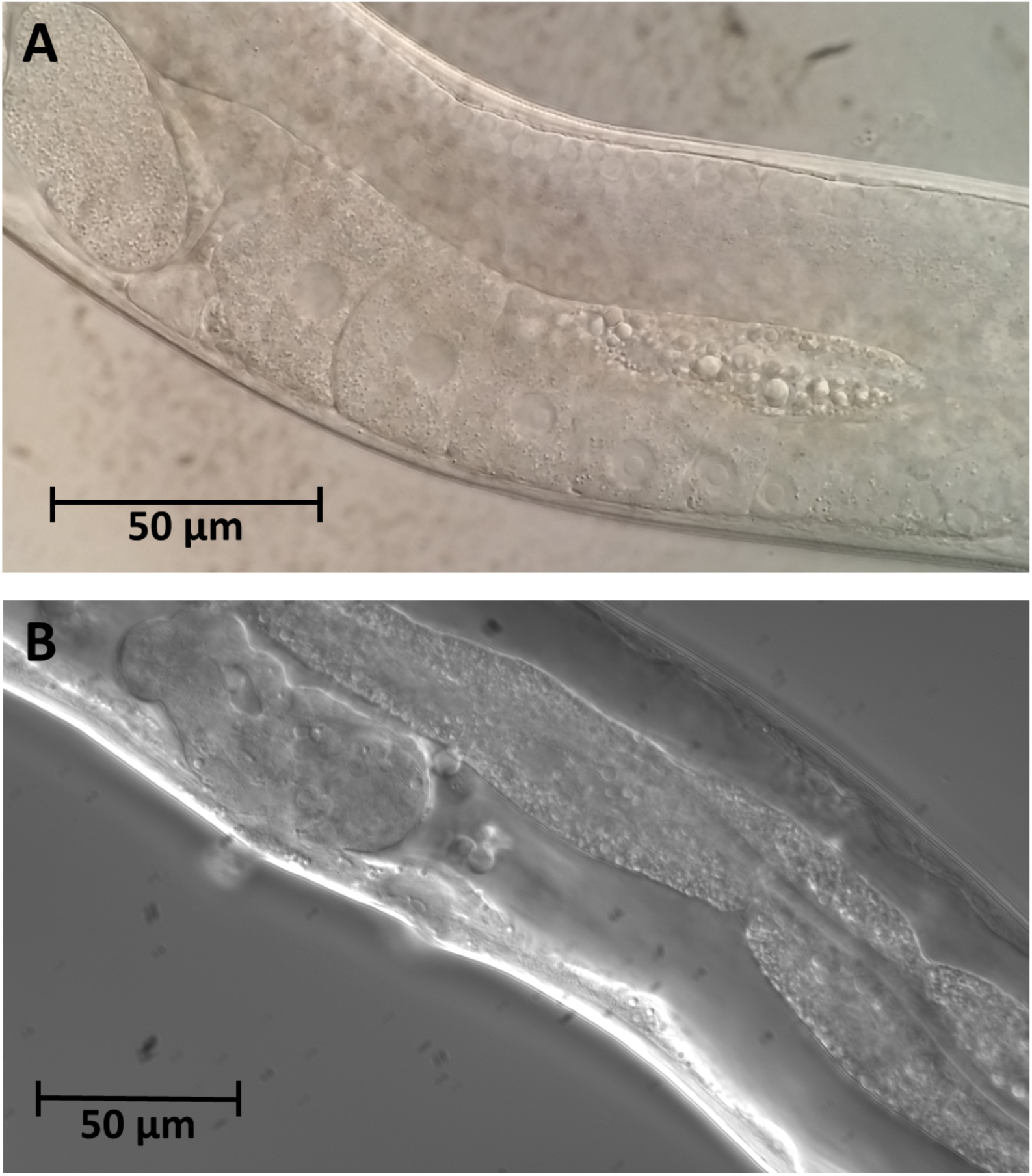
Loss of *adsl-1* severely disrupts gonad morphology. (**A**) An N2 adult animal showing normal gonad and oocyte development. (**B**) A representative *adsl-1(tm3228)* mutant adult has degenerate gonad morphology. No oocytes are present in the mutant animal.

**S2 Fig.**
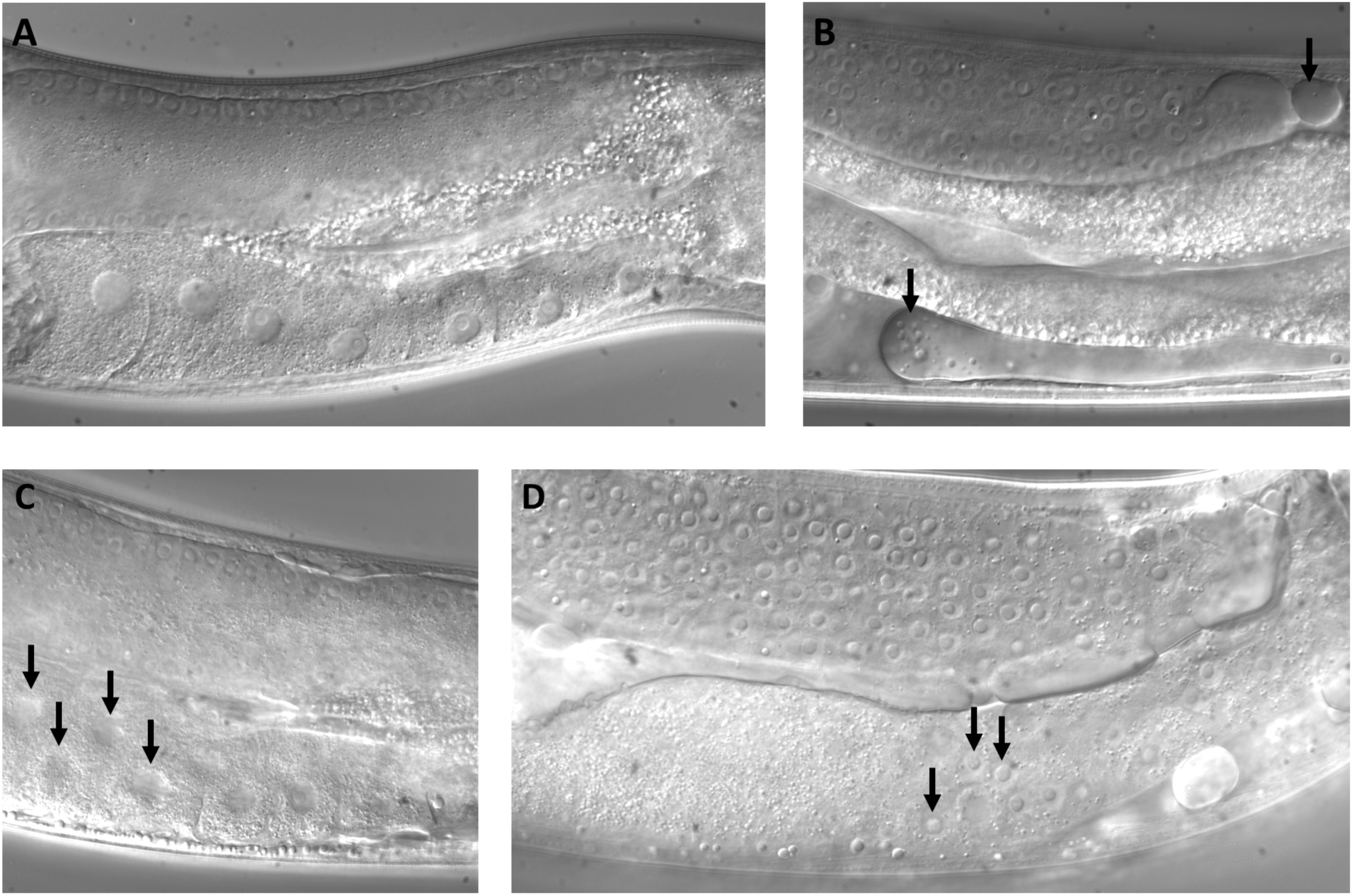
Knockdown of *adsl-1* causes defects in gonad morphology. (**A**) An *eri-1* adult animal with normal gonad development. (**B-D**) Adult *eri-1* animals exposed to *adsl-1*(RNAi) for 24 hours display a range of gonad morphology defects. Gonad deterioration (B), double oocytes in the proximal gonad (C), and germ cells in the proximal gonad (D) were all observed under these conditions of *adsl-1* knockdown.

## References

1. Jurecka A. Inborn errors of purine and pyrimidine metabolism. J Inherit Metab Dis. 2009;32:247–63.

2. Fang Y, French J, Zhao H, Benkovic S. G-protein-coupled receptor regulation of de novo purine biosynthesis: a novel druggable mechanism. Biotechnol Genet Eng Rev [Internet]. 2013;29:31–48. Available from: http://www.ncbi.nlm.nih.gov/pubmed/24568251

3. Jaeken J. AN INFANTILE AUTISTIC SYNDROME Succinyladenosine Case-reports. Pediatr Clin North Am. 1983;1058–61.

4. Stone RL, Aimi J, Barshop B a, Jaeken J, den Berghe G, Zalkin H, et al. A mutation in adenylosuccinate lyase associated with mental retardation and autistic features. Nat Genet. 1992;1(1):59–63.

5. Jaeken J, Wadman SK, Duran M, van Sprang FJ, Beemer FA, Holl RA, et al. Adenylosuccinase deficiency: an inborn error of purine nucleotide synthesis. Eur J Pediatr [Internet]. 1988 Nov [cited 2015 Jun 14];148(2):126–31. Available from: http://www.ncbi.nlm.nih.gov/pubmed/3234432

6. Zikanova M, Skopova V, Hnizda A, Krijt J, Kmoch S. Biochemical and structural analysis of 14 mutant ADSL enzyme complexes and correlation to phenotypic heterogeneity of adenylosuccinate lyase deficiency. Hum Mutat. 2010;31:445–55.

7. van den Bergh FA, Bosschaart AN, Hageman G, Duran M, Tien Poll-The B. Adenylosuccinase deficiency with neonatal onset severe epileptic seizures and sudden death. Neuropediatrics. 1998 Feb;29(1):51–3.

8. Mouchegh K, Zikánová M, Hoffmann GF, Kretzschmar B, Kühn T, Mildenberger E, et al. Lethal Fetal and Early Neonatal Presentation of Adenylosuccinate Lyase Deficiency: Observation of 6 Patients in 4 Families. J Pediatr. 2007;150.

9. Jurecka A, Zikanova M, Tylki-Szymanska A, Krijt J, Bogdanska A, Gradowska W, et al. Clinical, biochemical and molecular findings in seven Polish patients with adenylosuccinate lyase deficiency. Mol Genet Metab. 2008;94(4):435–42.

10. Gitiaux C, Ceballos-Picot I, Marie S, Valayannopoulos V, Rio M, Verrieres S, et al. Misleading behavioural phenotype with adenylosuccinate lyase deficiency. Eur J Hum Genet. 2009;17(1):133–6.

11. Krijt J, Kmoch S, Hartmannová H, Havlícek V, Sebesta I. Identification and determination of succinyladenosine in human cerebrospinal fluid. J Chromatogr B Biomed Sci Appl. 1999;726(1– 2):53–8.

12. Hürlimann HC, Laloo B, Simon-Kayser B, Saint-Marc C, Coulpier F, Lemoine S, et al. Physiological and toxic effects of purine intermediate 5-amino-4-imidazolecarboxamide ribonucleotide (AICAR) in yeast. J Biol Chem. 2011;286:30994–1002.

13. Van den Berghe G, Jaeken J. Adenylosuccinase deficiency. Adv Exp Med Biol [Internet]. 1986 Jan [cited 2015 Mar 16];195 Pt A:27–33. Available from: http://www.ncbi.nlm.nih.gov/pubmed/3014834

14. Van den Berghe G, Vincent MF, Jaeken J. Inborn errors of the purine nucleotide cycle:adenylosuccinase deficiency. J Inherit Metab Discov. 1997;20(2):193–202.

15. Ciardo F, Salerno C, Curatolo P. Neurologic aspects of adenylosuccinate lyase deficiency. J Child Neurol. 2001;16(5):301–8.

16. Jurecka A, Zikanova M, Kmoch S, Tylki-Szymańska A. Adenylosuccinate lyase deficiency. J Inherit Metab Dis [Internet]. 2015 Mar [cited 2015 Mar 15];38(2):231–42. Available from: http://www.pubmedcentral.nih.gov/articlerender.fcgi?artid=4341013&tool=pmcentrez&rendertype=abstract

17. Spiegel EK, Colman RF, Patterson D. Adenylosuccinate lyase deficiency. Mol Genet Metab. 2006;89(1–2):19–31.

18. Kuwabara PE, O'Neil N. The use of functional genomics in C. elegans for studying human development and disease. J Inherit Metab Dis. 2001;24:127–38.

19. Kappock TJ, Ealick SE, Stubbe J. Modular evolution of the purine biosynthetic pathway. Curr Opin Chem Biol [Internet]. 2000 Oct [cited 2015 Jun 14];4(5):567–72. Available from: http://www.ncbi.nlm.nih.gov/pubmed/11006546

20. White JG, Southgate E, Thomson JN, Brenner S. The Structure of the Nervous System of the Nematode Caenorhabditis elegans. Philos Trans R Soc Lond B. 1986;314(1165):1–340.

21. Gooding JR, Jensen M V., Dai X, Wenner BR, Lu D, Arumugam R, et al. Adenylosuccinate Is an Insulin Secretagogue Derived from Glucose-Induced Purine Metabolism. Cell Rep. The Authors; 2015;13(1):157–67.

22. Chen P, Wang D, Chen H, Zhou Z, He X. The non-essentiality of essential genes suggests a loss-of-function therapeutic strategy for loss-of-function human diseases. Genome Res. 2016;26:1355–62.

23. Ahringer J. Reverse genetics. In: The C. elegans Research Community, editor. WormBook [Internet]. 2006. Available from: http://www.wormbook.org

24. Duchaine TF, Wohlschlegel JA, Kennedy S, Bei Y, Conte D, Pang K, et al. Functional proteomics reveals the biochemical niche of C. elegans DCR-1 in multiple small-RNA-mediated pathways. Cell. 2006;124(2):343–54.

25. Ghosh R, Emmons SW. Episodic swimming behavior in the nematode C. elegans. J Exp Biol. 2008;211(Pt 23):3703–11.

26. Gray J. Undulatory propulsion in small organisms. Nature. 1951;168(4283):929–30.

27. Gray J, Lissmann HW. the Locomotion of Nematodes. J Exp Biol. 1964;41:135–54.

28. Butler VJ, Branicky R, Yemini E, Liewald JF, Gottschalk A, Kerr R a, et al. A consistent muscle activation strategy underlies crawling and swimming in Caenorhabditis elegans. J R Soc Interface. 2015;12(102):20140963.

29. Niebur E, Erdös P. Theory of the locomotion of nematodes: Dynamics of undulatory progression on a surface. Biophys J. 1991;60(5):1132–46.

30. Gao S, Zhen M. Action potentials drive body wall muscle contractions in Caenorhabditis elegans. Proc Natl Acad Sci U S A. 2011;108(6):2557–62.

31. Lewis JA, Wu C-H, Berg H, Levine JH. The genetics of levamisole resistance in the nematode Caenorhabditis elegans. Genetics. 1980;95:905–28.

32. Lewis JA, Wu C, Levine J, Berg H. Levamisole-resitant mutants of the nematode Caenorhabditis elegans appear to lack pharmacological acetylcholine receptors. Neuroscience. 1980;5(6):967–89.

33. Miller KG, Alfonso a, Nguyen M, Crowell J a, Johnson CD, Rand JB. A genetic selection for Caenorhabditis elegans synaptic transmission mutants. Proc Natl Acad Sci U S A. 1996;93(22):12593–8.

34. Mahoney TR, Luo S, Nonet ML. Analysis of synaptic transmission in Caenorhabditis elegans using an aldicarb-sensitivity assay. Nat Protoc. 2006;1(4):1772–7.

35. Allegra CJ, Hoang K, Yeh GC, Drake JC, Baram J. Evidence for direct inhibition of de novo purine synthesis in human MCF-7 breast cells as a principal mode of metabolic inhibition by methotrexate. J Biol Chem. 1987;262(28):13520–6.

36. Fairbanks LD, Rückemann K, Qiu Y, Hawrylowicz CM, Richards DF, Swaminathan R, et al. Methotrexate inhibits the first committed step of purine biosynthesis in mitogen-stimulated human T-lymphocytes: a metabolic basis for efficacy in rheumatoid arthritis? Biochem J. 1999;342(1):143–52.

37. Race V, Marie S, Vincent MF, Van den Berghe G. Clinical, biochemical and molecular genetic correlations in adenylosuccinate lyase deficiency. Hum Mol Genet. 2000;9(14):2159–65.

38. Karbowski J, Cronin CJ, Seah A, Mendel JE, Cleary D, Sternberg PW. Conservation rules, their breakdown, and optimality in Caenorhabditis sinusoidal locomotion. J Theor Biol. 2006;242(3):652–69.

39. Brenner S. The genetics of Caenorhabditis elegans. Genetics. 1974;77:71–94.

40. McKim KS, Peters K, Rose AM. Two types of sites required for meiotic chromosome pairing in Caenorhabditis elegans. Genetics. 1993;134(3):749–68.

41. Lu W, Clasquin MF, Melamud E, Amador-Noguez D, Caudy AA, Rabinowitz JD. Metabolomic analysis via reversed-phase ion-pairing liquid chromatography coupled to a stand alone orbitrap mass spectrometer. Anal Chem. 2010;82:3212–21.

